# Broad binaural fusion impairs segregation of speech based on voice pitch differences in a ‘cocktail party’ environment

**DOI:** 10.1101/805309

**Authors:** Yonghee Oh, Curtis L. Hartling, Nirmal Kumar Srinivasan, Morgan Eddolls, Anna C. Diedesch, Frederick J. Gallun, Lina A. J. Reiss

**Author notes:** Correspondence: Yonghee Oh, Department of Speech, Language, and Hearing Sciences, University of Florida, Gainesville, Florida 32610, USA.

## Abstract

In the normal auditory system, central auditory neurons are sharply tuned to the same frequency ranges for each ear. This precise tuning is mirrored behaviorally as the binaural fusion of tones evoking similar pitches across ears. In contrast, hearing-impaired listeners exhibit abnormally broad tuning of binaural pitch fusion, fusing sounds with pitches differing by up to 3-4 octaves across ears into a single object. Here we present evidence that such broad fusion may similarly impair the segregation and recognition of speech based on voice pitch differences in a ‘cocktail party’ environment. Speech recognition performance in a multi-talker environment was measured in four groups of adult subjects: normal-hearing (NH) listeners and hearing-impaired listeners with bilateral hearing aids (HAs), bimodal cochlear implant (CI) worn with a contralateral HA, or bilateral CIs. Performance was measured as the threshold target-to-masker ratio needed to understand a target talker in the presence of masker talkers either co-located or symmetrically spatially separated from the target. Binaural pitch fusion was also measured. Voice pitch differences between target and masker talkers improved speech recognition performance for the NH, bilateral HA, and bimodal CI groups, but not the bilateral CI group. Spatial separation only improved performance for the NH group, indicating an inability of the hearing-impaired groups to benefit from spatial release from masking. A moderate to strong negative correlation was observed between the benefit from voice pitch differences and the breadth of binaural pitch fusion in all groups except the bilateral CI group in the co-located spatial condition. Hence, tuning of binaural pitch fusion predicts the ability to segregate voices based on pitch when acoustic cues are available. The findings suggest that obligatory binaural fusion, with a concomitant loss of information from individual streams, may occur at a level of processing before auditory object formation and segregation.

## INTRODUCTION

For normal-hearing (NH) listeners, the two ears provide essentially matched spectral information for a given signal. Binaural pitch fusion occurs only for tones of similar pitch, with tolerance of differences for dichotically presented tones to be less than 0.1-0.2 octaves (Thurlow and Bernstein, 1957; Odenthal, 1963; van den Brink, 1976). This sharp binaural tuning is mirrored in central auditory neurons which have sharp, matched bilateral frequency tuning in the medial superior olive and inferior colliculus, but not necessarily in auditory cortex (e.g. Pena et al., 2001; McLaughlin et al., 2007; Sollini et al., 2017).

In contrast, hearing-impaired listeners with hearing aids (HAs) and/or cochlear implants (CIs) exhibit abnormally broad binaural pitch fusion, fusing tones (or electrodes) with pitches differing by up to 3-4 octaves across ears into a single percept (Reiss et al., 2014; 2017; 2018). Further, this broad fusion leads to the weighted averaging of different pitches across the ears (Reiss et al., 2014; Oh and Reiss, 2017), as well as binaural interference for spectral shape perception of vowel stimuli (Reiss et al., 2016). It is not yet clear where abnormal integration manifests in the auditory system, or how it would arise; one possible cause is reduced pruning and tonotopic refinement during development, as suggested by an association of broad fusion with childhood onset of hearing loss (Reiss et al., 2017).

Broad binaural pitch fusion could interfere with the segregation of auditory objects based on pitch differences, such as multiple voices in a multi-talker listening environment. In particular, speech perception in noisy listening environments requires the ability to separate this target speech from interfering concurrent sounds (maskers) and selectively attend to the speech source of interest (target). A multi-talker listening situation is particularly difficult because the target and competing masker speech sources are acoustically similar in spectral shape and temporal envelope. This multi-talker listening situation is well known as the “cocktail party effect” (Cherry, 1953).

For NH listeners, performance in multi-talker environments can be improved by various acoustic factors. One major factor, spatial separation of the target from the maskers, can enhance a listener’s speech perception ability compared to the co-located target-masker configuration. This improvement is referred to as spatial release from masking (SRM). In NH listeners, the amount of SRM can be as large as 18 dB depending on spatial separation in the horizontal plane (Arbogast et al., 2002). Another factor, a gender difference between the target and the masker (e.g., male target and female masker or vice versa), can lead to a voice pitch difference cue which also improves the ability to perceive the target speech (e.g. Brungart, 2001). In this study, we will refer to this benefit from voice pitch differences as voice gender release from masking (VGRM), often previously referred to as “informational masking” (e.g. Brungart, 2001; Arbogast et al., 2002).

In HI listeners, binaural device use, including HAs or CIs in both ears (bilateral HAs or bilateral CIs) or a CI with a contralateral HA (bimodal CI+HA), can be a major factor for binaural listening advantages in both spatial and voice gender difference conditions (Litovsky et al., 2006; Marrone et al., 2008; Visram et al., 2012; Bernstein et al., 2016). However, benefits from binaural devices are highly variable, and often provide little speech perception benefit or even interfere with speech perception, compared to monaural device use (Litovsky et al., 2006; Ching et al., 2007; Reiss et al., 2016). Variation in binaural pitch fusion may explain the variability in benefits seen in the previous studies.

The goal of this study was to investigate how binaural pitch fusion affects the ability to benefit from voice pitch differences for separation of voices in a realistic multi-talker environment. Binaural pitch fusion ranges and speech recognition thresholds with multi-talker maskers were measured in NH listeners and hearing-impaired (HI) listeners groups including bilateral HA users, bimodal CI+HA users, and bilateral CI users. The results were analyzed to determine the benefit from voice pitch and spatial cues, and the relationship between the breadth of binaural pitch fusion and overall performance on speech perception in background talkers.

## MATERIALS AND METHODS

### Participants

These studies were conducted according to the guidelines for the protection of human subjects as set forth by the Institutional Review Boards (IRBs) of both Oregon Health & Sciences University and the Portland VA Medical Center, and the methods employed were approved by these IRBs. Forty-seven adult subjects, consisting of eleven NH listeners ranging in age from 36 to 67 years (mean and standard deviation (std) = 50.0 ± 9.9 years; 4 males and 7 females), twelve bilateral HA users ranging in age from 30 to 75 years (mean and std = 53.8 ± 16.7 years; 2 males and 10 females), twelve bimodal CI+HA users ranging in age from 30 to 80 years (mean and std = 56.2 ± 14.6 years; 5 males and 7 females), and twelve bilateral CI users ranging in age from 20 to 66 years (mean and std =53.0 ± 14.8 years; 7 males and 5 females), participated in this study. A Kruskal-Wallis H test showed that there were no significant age differences between these four listener groups (H(3)=1.817, p=0.611). All subjects were screened for normal cognitive function using the 10-minute Mini Mental Status Examination (MMSE) with a minimum score of 27 out of 30 required to qualify (Folstein et al., 1975; Souza et al., 2007). It should be noted that each subject population in this study was purposely sampled in accordance with the study design which requires a range of binaural fusion from narrow to broad to investigate correlations over this range. In particular, the sampling of the NH subject population was highly biased because many NH subjects with narrow fusion were excluded, and is not representative of the true population mean.

Normal hearing was defined as air conduction thresholds ≤ 25 dB hearing level (HL) from 125 to 4000 Hz. Mean pure-tone averages at octave interval frequencies between 125 and 4000 Hz for NH subjects were 12.6 ± 2.2 dB HL for the left ear and 11.5 ± 1.4 dB HL for the right ear. Bilateral HA users had moderate to severe hearing losses in both ears and relatively symmetric losses between ears, with the exception of subject H1. Mean pure-tone averages were 56.5 ± 10.8 and 57.7 ± 10.5 dB HL for left and right ears, respectively. For bimodal CI+HA users, mean pure-tone averages were 81.2 ± 17.8 dB HL in the non-implanted ear. Note that all of the bimodal CI+HA users in the study used a contralateral HA together with the CI on a regular basis. Both bilateral HA and bimodal CI+HA users had unaided residual hearing (<110 dB HL) up to 2000 Hz in their both ears and in their non-implanted ear, respectively. Figure 1A shows group-averaged audiograms for NH subjects (thick solid lines) and individual audiograms for bilateral HA subjects (lines with open symbols) for left and right ears. Figure 1B shows unaided individual audiograms in the non-implanted ear for bimodal CI+HA subjects.

**Figure 1.**
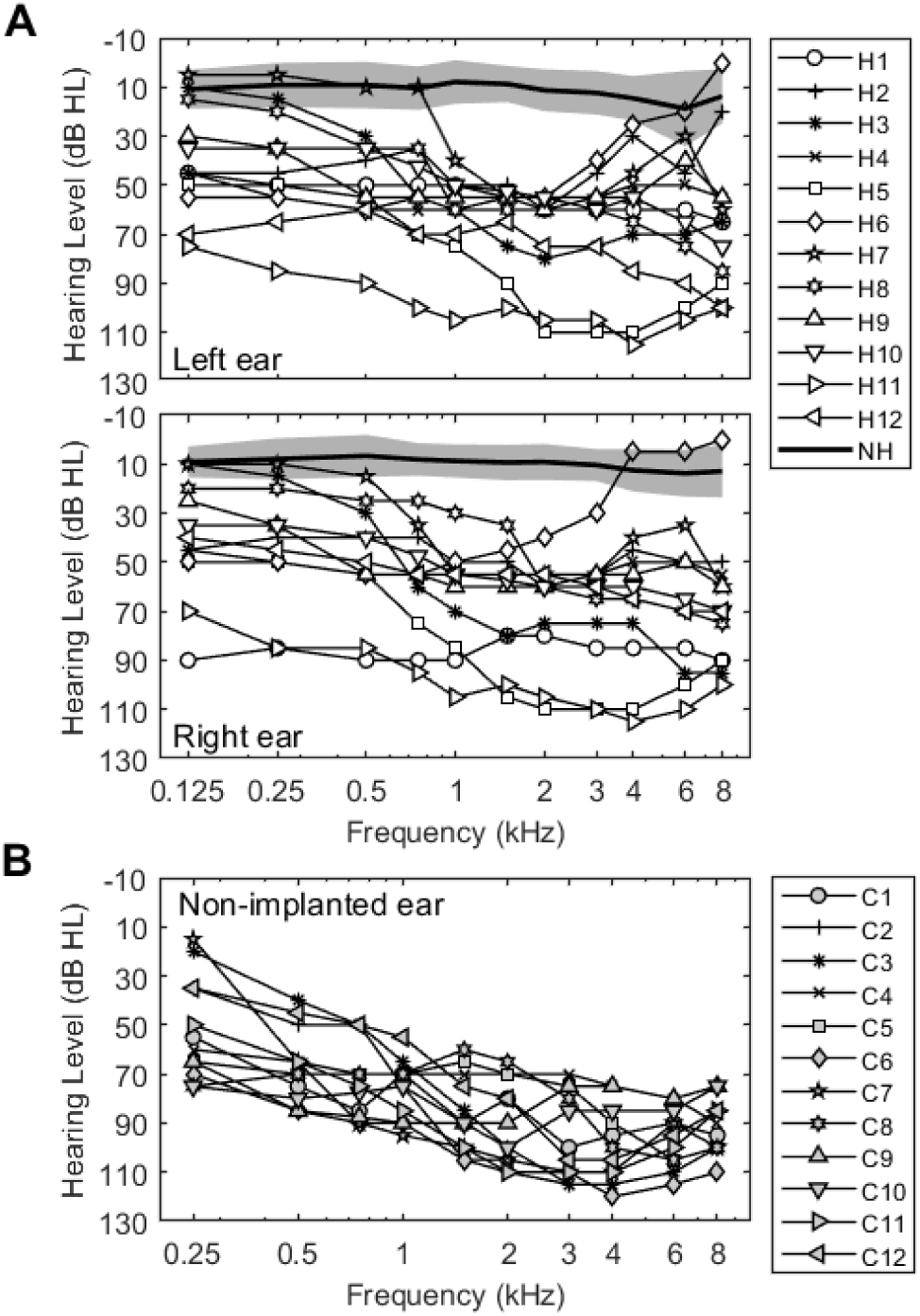
**A.** Unaided audiograms for the NH and bilateral HA subjects in this study. Solid thin lines show individual thresholds for bilateral HA users. Solid thick lines and shaded areas represent averaged thresholds and standard deviations for NH subjects. **B.** Individual unaided audiograms for non-implanted ears of the bimodal CI+HA users.

All HI subjects were required to have at least one year of experience with their hearing devices, and have monaural word intelligibility scores of 65% or higher on the Consonant Nucleus Consonant (CNC) word test with both devices. All HAs for both bilateral HA and bimodal CI+HA subjects were verified to meet NAL-NL2 (National Acoustics Laboratory, Australia) targets (speech stimuli at 50, 65, and 75 dB SPL) using real-ear measurements in order to provide suitable amplification for a subject’s hearing loss. In all subject groups, except bilateral CI users, tympanometry was also conducted to verify normal middle ear function. Demographic information for all CI subjects including age, gender, duration of bimodal, duration of CI use for each ear, stimulation pulse width, and CI external and internal (and/or HA) model(s) are shown in Table 1.

**Table 1.** Demographic information for cochlear implant (CI) users: age, duration of bimodal, duration of CI use (left; right for bilateral CI subjects), HA and CI (external / internal) model, pulse-width, and reference ear used in testing. Sub = subject, Dur = duration, Ref = reference, Std = Standard; R=right; L=left, and n/a=not applicable. CI24R, CI24RE, and CI512 are all modiolar arrays. CI24M, CI422, Flex28, and Standard are all straight lateral wall arrays.

| Sub.<br>ID # | Sex | Dur.<br>bimodal<br>(years) | Dur.<br>CI use<br>(years,<br>L;R if<br>diff.) | HA/ CI Brand and Model<br>(CI, external/internal) |  | Stimulus<br>Pulse-width<br>(μs, L;R if<br>diff.) | Ref.<br>Ear |
| --- | --- | --- | --- | --- | --- | --- | --- |
|  |  |  |  | Left | Right |  |  |
| Bimodal CI+HA (N=12) |  |  |  |  |  |  |  |
| CI1 | F | 7 | 7 | Cochlear N6 / CI24RE | Widex Diva | 25 | L |
| CI2 | M | 1 | 1 | Widex Dream | Cochlear N6 / CI422 | 25 | R |
| CI3 | F | 2 | 4 | Oticon Adapto | MedEL Opus2 / Std Sonata | 25.42 | R |
| CI4 | F | 4 | 4 | Resound BTE | Cochlear N6 / CI422 | 25 | R |
| CI5 | M | 3.5 | 3.5 | Cochlear N6 / CI422 | Resound Linx2 | 25 | L |
| CI6 | M | 8 | 8 | Resound Enzo5 | Cochlear N6 / CI24RE | 25 | R |
| CI7 | F | 3 | 5 | Cochlear N5 / CI422 | Unitorn 360+ | 25 | L |
| CI8 | F | 4 | 4 | Resound Linx2 | Cochlear N5 / CI512 | 25 | R |
| CI9 | F | 6 | 6 | Cochlear N5 / CI512 | Phonak Naida | 25 | L |
| CI10 | M | 4 | 4 | Phonak Naida | Cochlear N5 / CI512 | 37 | R |
| CI11 | M | 5 | 5 | Phonak Naida | Cochlear N5 /CI24RE | 25 | R |
| CI12 | F | 4 | 4 | Phonak Naida | Cochlear N5 / CI24RE | 25 | R |
| Bilateral CI (N=12) |  |  |  |  |  |  |  |
| BI1 | M | 1.5 | 3;4.5 | Cochlear N5 / CI512 | Cochlear N5 / CI24RE | 25 | R |
| BI2 | M | 1 | 7;7 | Cochlear N6 / CI24R | Cochlear N6 / CI24RE | 25 | L |
| BI3 | M | 2.3 | 1;4.1 | MedEL Opus2 / Flex 28 | MedEL Opus2 / Std Concerto | 35;28.33 | L |
| BI4 | M | 0 | 11;1.5 | Cochlear N6 / CI24R | Cochlear N6 / CI422 | 25 | R |
| BI5 | F | 0 | 13;6.7 | Cochlear N5 / CI24R | Cochlear N5 / CI24RE | 25 | L |
| BI6 | F | 0 | 1.7;1.3 | Cochlear N6 / CI24RE | Cochlear N6 / CI24RE | 25 | R |
| BI7 | M | 7 | 15;5.9 | MedEL Opus2 / Std C40+ | MedEL Opus2 / Std Sonata | 26.67;25.83 | L |
| BI8 | M | 3.5 | 2;3.5 | Cochlear N6 / CI522 | Cochlear N6 / CI422 | 25 | R |
| BI9 | F | 0 | 4.6;3.6 | MedEL Opus2 / Std Concerto | MedEL Opus2 / Std Concerto | 30.83 | R |
| BI10 | F | 1 | 2;15 | Cochlear N6 / CI422 | Cochlear N6 / CI24R | 25 | L |
| BI11 | F | 0.5 | 1.7;2.2 | Cochlear N5 / CI24RE | Cochlear N5 / CI24RE | 25 | L |
| BI12 | M | 1.6 | 11;17 | Cochlear N6 / CI24R | Cochlear N6 / CI24M | 37 | L |

### Stimuli and Procedures

Two main experiments were conducted in this study: binaural pitch fusion range measurement and speech recognition in speech threshold measurement. The results for all experiments were averaged with three separate runs for each condition. All statistical analyses were conducted in SPSS (version 25, IBM).

#### Binaural pitch fusion range measurement

Binaural pitch fusion experiments were conducted in a double–walled, sound attenuated booth. All stimuli were generated via computer to control and synchronize electric and/or acoustic stimulus presentation.

Acoustic stimuli for both ears in NH listeners and bilateral HA users, and the non-implanted ear in bimodal CI+HA users were digitally generated at a sampling rate of 44.1 kHz with C++ or MATLAB, delivered using an ESI Juli sound card, TDT PA5 digital attenuator and HB7 head phone buffer, and presented over Sennheiser HD-25 headphones. Headphone frequency responses were equalized using calibration measurements obtained with a Brüel & Kjær sound level meter with a 1-inch microphone in an artificial ear. Acoustic stimuli consisted of 1500-ms harmonic tone complexes for both NH and bilateral HA subjects and 1500-ms sinusoidal pure tones for bimodal CI+HA subjects, with 10-ms raised-cosine onset/offset ramps. Previous studies of binaural fusion had used pure tones for all subject groups, which limited the frequencies that could be tested in NH and bilateral HA groups to above the upper binaural beat limit frequency of ∼1500 Hz (Perrott & Nelson, 1969); tones below 1500 Hz elicit binaural beats which could interfere with fusion judgements. In this study, harmonic tone complexes were used instead of pure tones for NH and HA subjects in order to measure fusion ranges in the lower frequency ranges relevant for voice pitch discrimination, without eliciting binaural beats. Pure tones were still used for bimodal CI+HA subjects, because binaural beats do not occur in this group.

For CI subjects, electric stimuli for the CI ear were delivered directly to the CI using custom research software and hardware for each company. For Cochlear, NIC2 software interface was used with L34 processors and programming pods. For MED-EL, RIB2 software interface and National Instruments PCIe-6351 card were used to control stimulation through a research interface box with dual outputs. Each stimulus consisted of biphasic pulse trains presented at 1200 pulses per second, using a monopolar ground, with pulse widths ranging from 25 to 37-µs depending on the individual subject’s clinical mapping.

Prior to the binaural fusion range measurements, loudness balancing was conducted sequentially across frequencies or electrodes and across ears using a method of adjustment. For NH listener and bilateral HA user groups, 300-msec tones at 0.125, 0.25, 0.375, 0.5, 0.625, 0.75, 0.875, 1, 1.25, 1.5, 2, 3, and 4 kHz in the reference ear were initialized to “medium loud and comfortable” levels corresponding to a 6 or “most comfortable” on a visual loudness scale from 0 (no sound) to 10 (too loud). Loudness for the comparison ear was then adjusted for each frequency to be equally loud to a tone in the reference ear during sequential presentation across the ears, based on subject feedback. Here, all loudness balancing adjustments were repeated with a fine attenuation resolution (0.1 dB steps for bilateral HA and 0.5 dB steps for NH listeners) until equal loudness was achieved with all comparison sequences within and across ears. The averaged comfortable sound levels were 65 ± 4 / 65 ± 4.1 dB sound pressure level, SPL (left / right ear) for NH listeners and 90 ± 1.4 / 91 ± 1.7 dB SPL (left / right ear) for bilateral HA users.

For both CI user groups, the level of the electric stimulation for each electrode was similarly loudness balanced as a “medium loud and comfortable” current level. Then, for the bimodal CI+HA users, acoustic tones were presented to the contralateral, non-implanted ear and set to “medium loud and comfortable” levels again using the same loudness scale as for electric levels. The averaged comfortable sound levels were 95 ± 4.2 dB SPL for the non-implanted ear. Tone frequencies that could not be presented loud enough to be considered a “medium loud and comfortable” level were excluded because of the limited range of residual hearing in the contralateral ear. For the bilateral CI users, the contralateral CI electrode was loudness-balanced sequentially with the reference electrodes.

The electrodes/frequencies and order of presentation were randomized to minimize the effect of biases such as time-order error and underestimation or overestimation of the loudness (Florentine et al., 2011). This loudness balancing procedure was performed to minimize use of level-difference cues and maximize focus on pitch differences as the decision criteria.

Binaural pitch fusion range measurements were then performed to measure the fusion ranges over which dichotic pitches were fused. With a1500-msec duration, the following stimulus types were used in different listener groups: 1) dichotic harmonic tone complexes in NH and bilateral HA subjects; 2) dichotic pure tone and electric pulse train in bimodal CI+HA subjects; and 3) dichotic electric pulse trains in bilateral CI subjects. For all fusion range measurements, a method of constant stimuli procedure was used. The reference ear and stimulus was fixed in the designated “reference ear”, and the contralateral, comparison ear stimulus was varied across trials. For NH listeners, the reference ear was randomized. For bimodal CI+HA listeners, the CI ear was the reference ear. For bilateral HA and bilateral CI groups, the ear with worse within-ear frequency or electrode discrimination (data not shown) was assigned to be the reference ear so that the resolution of comparison stimulus testing would be maximized using the better ear, instead of limited by the worse ear.

Fusion range procedure varied depending on group. First, for NH and bilateral HA subjects, harmonic tone complexes with the reference fundamental frequency (F_0ref_) fixed at 200 Hz was used. The comparison stimuli consisted of other harmonic complexes with fundamental frequencies (F_0comp_) sampled around the reference with 1/64 to 1/16 octave steps and varied pseudo-randomly across trials. The number of harmonic components were fixed at four.

Second, for the fusion range measurement in bimodal CI+HA subjects, the reference stimulus (electrode 12, 18, or 20 for Cochlear and 5, 7, or 9 for MEL-EL) was held constant in one ear, and the comparison stimuli were pure tones of various frequencies sampled in 1/32 to 1/4 octave steps around the reference and varied pseudo-randomly across trials.

Third, for the fusion range measurement in bilateral CI subjects, the reference stimulus (same reference electrodes as for bimodal CI+HA subjects) was held constant in one ear, and the comparison electrodes (even-numbered electrodes for Cochlear and all electrodes for MED-EL) were varied pseudorandomly across trials.

Subjects were asked to indicate whether they heard a single fused sound or two different sounds. If a single sound was heard, subjects were instructed to indicate whether they heard that sound as a single fused sound (“Same”). If two different sounds were heard, subjects were instructed to indicate which ear had the higher pitch (“Left higher” or “Right higher”) as a check of whether two sounds were really heard. A “Repeat” button was also provided to allow subjects to listen to the stimuli again. Responses were obtained using a touch screen monitor.

Fusion functions were computed as the averages of the subject responses to the multiple (six to seven) presentations of each reference and comparison stimulus pair and expressed as a function of comparison tone frequency (NH listeners, bilateral HA users, and bimodal CI+HA users) or electrode (bilateral CI users). Values near 0 indicate comparison stimuli that did not often fuse with the reference stimulus (were heard as two sounds), while values near 1 indicate comparison stimuli that were often fused with the reference stimulus (were heard as one sound). Example fusion functions for a representative subject in each subject group are shown in the insets in Figure 2, as solid lines with circle symbols. Vertical dotted lines indicate 50% points on the fusion function, and the fusion range was defined as the range between these two lines (horizontal arrows in Fig. 2), i.e. frequencies or electrodes were fused more than 50% of the time. Fusion range is thus a measure of the breadth of fusion.

**Figure 2.**
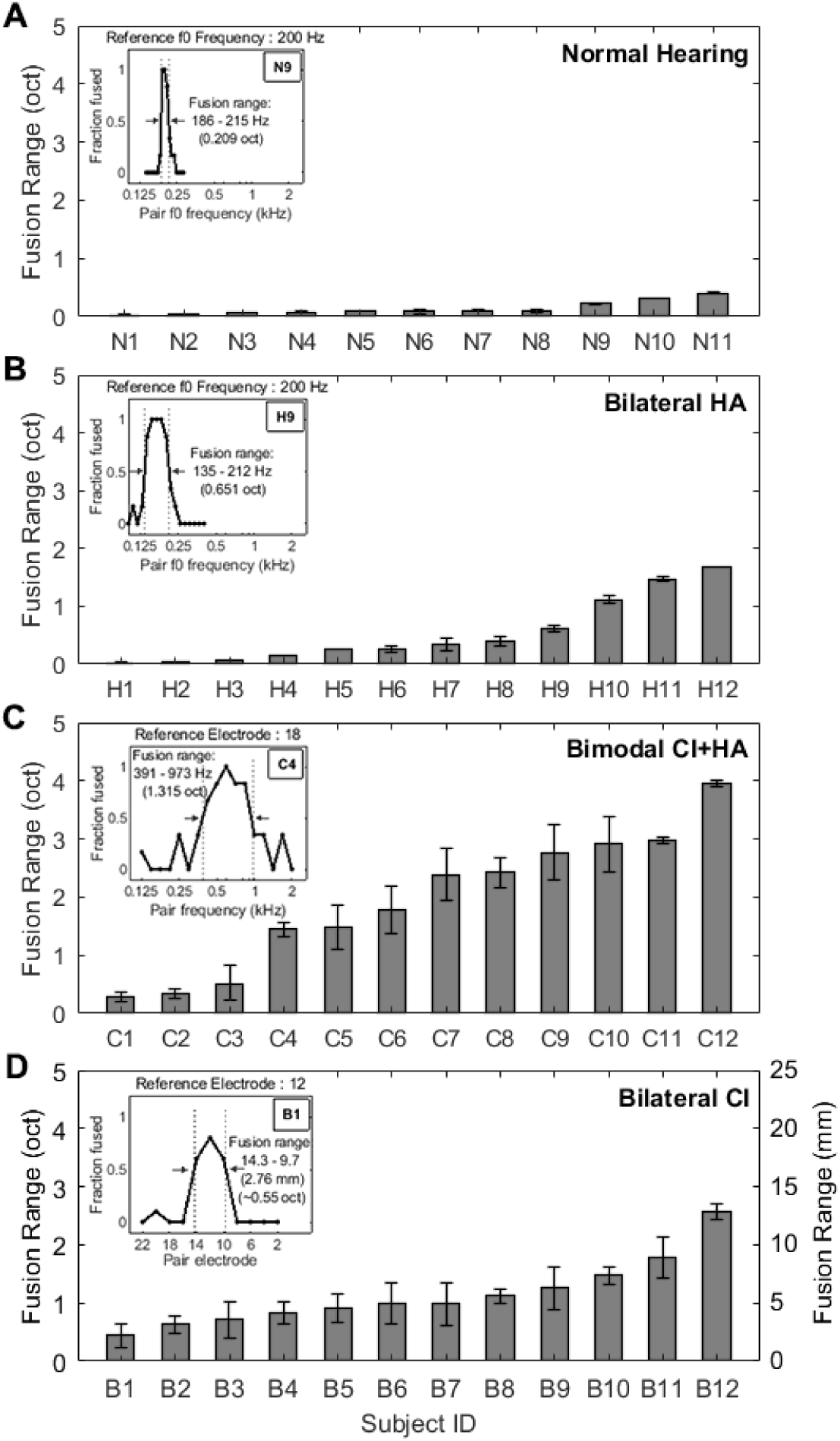
Individual fusion range results by group. **A**) NH listeners; **B**) bilateral HA users; **C**) bimodal CI+HA users; **D**) bilateral CI users. Error bars represent standard deviations around the mean. A sample fusion function inset within each panel illustrates the fusion ranges of the 50% points (vertical dotted lines) on the fusion function. In panel D, the scale of the bilateral CI data, in mm, is approximated to 1 octave per 5 mm (Greenwood, 1990), and scaled accordingly for comparison with the NH, bilateral HA and bimodal CI+HA data.

For the bilateral CI users, to allow comparisons across CI electrode arrays with different inter-electrode distances, results in units of electrodes were converted into mm (measured between the centers of adjacent electrodes) using the following inter-electrode distances: 1.9 and 2.4 mm for MED-EL Flex 28 and Standard arrays, 0.75 mm for Cochlear CI24M, and variable distances between 0.4-0.81 mm for Cochlear CI24R/CI24RE/CI512 and between 0.85-0.95 mm for Cochlear CI422 arrays.

#### Measurement of speech recognition threshold in a multi-talker environment

Speech recognition experiments were conducted in the anechoic chamber located at the National Center for Rehabilitative Auditory Research (NCRAR). All bilateral HA subjects used lab loaner HA devices (Phonak/Ambra), and all CI users used their personal HA and/or CI devices with disabling all extra processing features for hearing devices including adaptive/automatic gain control, frequency lowering, directional microphones, and noise reduction. All subject groups were tested binaurally. A subset of NH and bilateral HA subjects were further tested in one or both monaural listening conditions with the non-test ear plugged and muffed.

All speech stimuli were digitally processed in MATLAB to have a sampling rate of 44.1 kHz. Stimuli were presented through a bank of three eight-channel amplifiers (Ashlys/ne4250) and 24 frequency-equalized loudspeakers calibrated by a Brüel & Kjær sound level meter. The loudspeakers were arranged in a circle in the horizontal plane with 15° increments surrounding the listener and equidistant at 2 m from the listener’s head.

All speech stimuli were drawn from the Coordinate Response Measure (CRM; Bolia et al., 2000) speech corpus, which consists of sentences in the form “*Ready* [*call sign*] *go to* [*color*] [*number*] *now*.” In this study, speech stimuli were presented with a 20% slower speaking rate than the original CRM corpus stimuli because some HI users had difficulties in understanding target-only stimuli at the original speaking rate. A custom MATLAB implementation of a modified pitch synchronous overlap add (PSOLA) technique (Moulines and Laroche, 1995) was used to expand CRM sentences by 20%. There are eight possible *call signs* (Arrow, Baron, Charlie, Eagle, Hopper, Laker, Ringo, and Tiger), and 12 keywords: four *colors* (red, green, white, and blue) and the *numbers* (1–8). All possible combinations of the *call signs*, *colors*, and *numbers* were spoken by four male (F_0_ = 100 ± 7 Hz) and four female talkers (F_0_ = 204 ± 12 Hz). Note that fundamental frequency (F_0_), which represents the voice pitch, was estimated using the cepstrum algorithm in MATLAB where the output is the Fourier transform of the log of the magnitude spectrum of the input waveform (Flanagan, 1965). F_0_ for each talker was averaged across all of that talker’s CRM speech stimuli.

Target speech stimuli were presented from directly in front of the listener with a fixed sound presentation level of 60 dB SPL. Masker speech stimuli were presented in one of two spatial configurations: co-located (target at 0°, maskers at 0°) or 60° symmetrical separations (target at 0°, maskers at ±60°). Only symmetrical target-masker separation conditions were considered in order to minimize availability of the better ear cue (monaural head shadow effect; Shaw, 1974; Kidd et al., 1998) and maximize reliance on spatial cues or voice pitch cues for source segregation.

These two spatial conditions were tested with four different gender combinations of target and maskers: MM (male target, male maskers), MF (male target, female maskers), FF (female target, female maskers), and FM (female target, male maskers), for a total of 2 × 4 = 8 conditions. In each trial, the subject was instructed to face the front speaker and attend to the target sentence, always identified here by the *call sign* “Charlie”, and indicate the target *color* and *number* keywords from the 32 possible *color*/*number* combinations. The masker sentences had exactly the same form as the target but a different *call sign*, *color*, and *number*, randomly selected on each trial. The target and masker sentences were randomized from eight talkers (4 males and 4 females) for each target-masker gender combination at each trial, and they were temporally aligned at the beginning and were roughly the same total duration.

Responses were obtained using a touch screen monitor located on a stand within arm’s reach of the listener seated in the middle of the anechoic chamber. The monitor was directly in front of the listener but below the plane of the loudspeakers. Feedback was given after each presentation in the form of “Correct” or “Incorrect”. Approximately one second of silence followed the response being registered, prior to the next stimulus presentation.

The masker sound presentation level was adaptively varied at each trial to find the target-to-masker ratio (TMR), or the masker level yielding 50% correct recognition of the target *color* and *number* (i.e. 1/32 chance), using a one-up/one-down procedure (Levitt, 1971). The initial level for the masker sentence was set at 30 dB SPL and increased in level by 5 dB for each correct response until an incorrect response occurred, then decreased in level for each incorrect response until a correct response, and so on. This was repeated until three reversals in direction were obtained, at which point the step size was changed to 1 dB and six more reversals were measured. The TMR was estimated as the average of the last six reversals.

The order of conditions was randomized. The TMR was averaged over three tracks for each condition. Note that TMR indicates the difference in level between the target and each masker in the symmetrical target-masker separation conditions, while signal-to-noise ratio (SNR) refers to difference between the target and the combined masker level. For example, if the target level is 60 dB SPL and each masker is also 60 dB SPL, the TMR would be 0 dB, and the overall SNR would be approximately −3 dB.

## RESULTS

### Binaural pitch fusion differences in NH, HA, and CI listeners

Figure 2 shows individual fusion range results for NH (Fig. 2A), bilateral HA (Fig. 2B), bimodal CI+HA (Fig. 2C), and bilateral CI listeners (Fig. 2D). Note that stimuli differed across groups, with harmonic tone complexes used for NH and HA groups, and pure tones or single electrodes used for both CI groups. Fusion ranges are shown as gray bars in ascending order of magnitude. The NH subjects exhibited narrow fusion ranges (0.03 to 0.40 octaves with mean and std = 0.14 ± 0.12 octaves), while HI groups showed relatively broader fusion ranges (bilateral HA users: 0.03 to 1.67 octaves with mean and std = 0.53 ± 0.57 octaves; bimodal CI+HA users: 0.28 to 3.95 octaves with mean and std = 1.98 ± 1.19 octaves; bilateral CI users: 2.19 to 12.8 mm with mean and std = 5.71 ± 2.90 mm). The average fusion range in mm for bilateral CI users corresponds to slightly over an acoustic octave (∼1.11± 0.63 octaves, see the right y-axis labels scaled in octaves in Fig. 2D) in the human ear where 5 mm is approximately 1 octave (in the 1-8 kHz range likely to be stimulated by the electrodes; Greenwood, 1990).

Note that the population mean of the fusion range was not compared because different stimuli were used in different listener groups (NH and HA user groups: dichotic harmonic tone complexes; both CI user groups: pure tones or single electrodes). In addition, the subjects in each listener group were purposely sampled with a narrow to broad fusion range so that the samples didn’t reflect the true population distribution of the fusion range.

In both CI user groups, the fusion range results were averaged across all reference stimuli (electrode 12, 18, or 20 for Cochlear and 5, 7, or 9 for MEL-EL). When the fusion ranges were compared among these reference stimuli, no significant differences were seen within CI subject groups (F_2,66_=0.389, p=0.679, two-way ANOVA).

### Effects of spatial separation and voice pitch differences on speech recognition in noise

The top row of Figure 3 shows the group mean TMR thresholds (±1 standard deviation around the mean) for the 8 different conditions (4 target-masker gender combinations × 2 spatial separations) for NH (Fig. 3A), bilateral HA (Fig. 3B), bimodal CI+HA (Fig. 3C), and bilateral CI listeners (Fig. 3D). Note that the TMR thresholds of the two same-gender conditions (MM and FF) were similar at each spatial configuration across all listener groups, as were those of the two different-gender conditions (MF and FM). In order to more clearly show the effects of same-gender versus different-gender maskers, the bottom row of Figure 3 shows the TMR thresholds averaged over these similar same-gender (MM and FF) versus different-gender conditions (MF and FM) for each spatial condition. Generally, smaller or more negative TMR thresholds indicate better speech recognition ability in noise.

**Figure 3.**
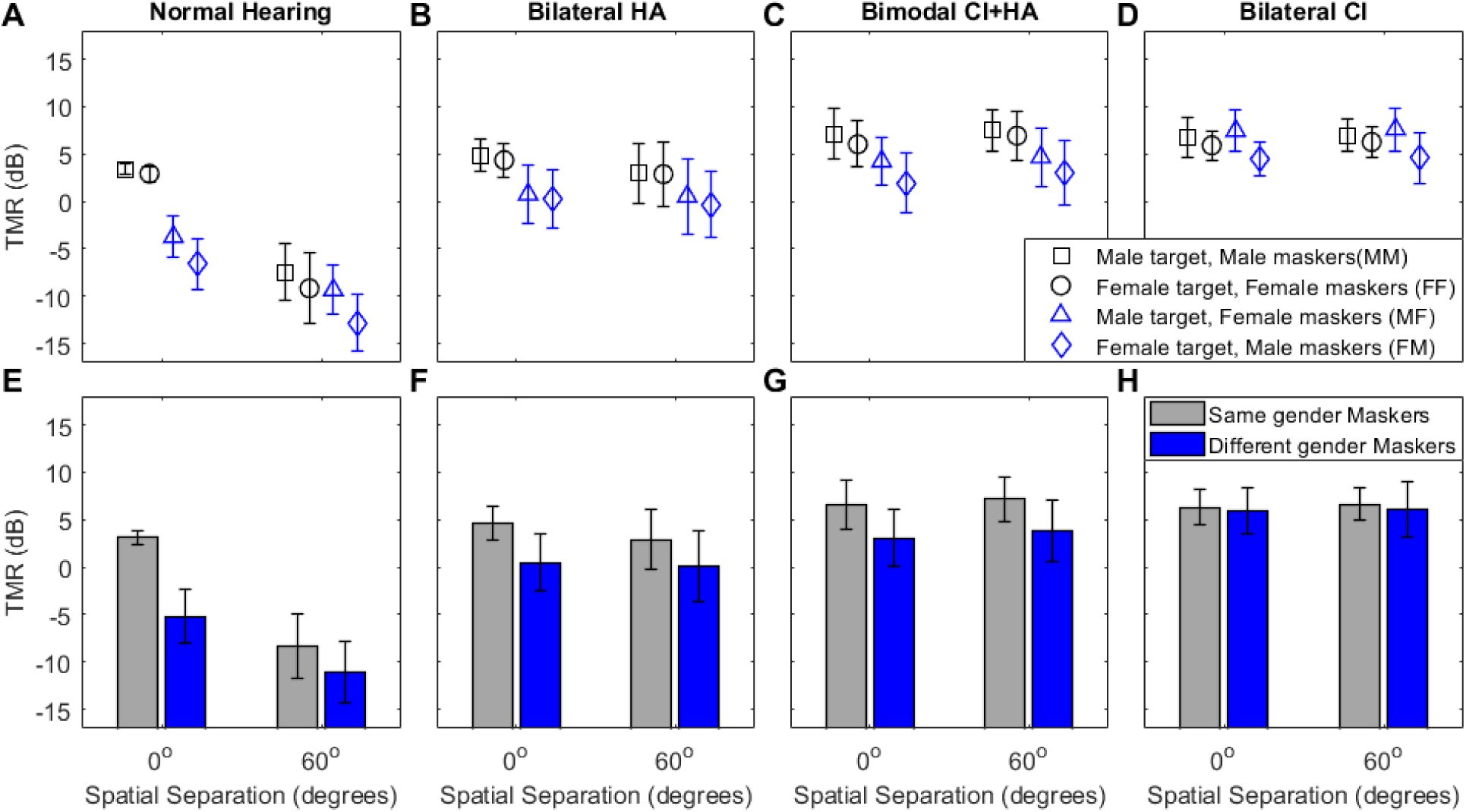
Average target to masker ratio (TMR) plotted by target-masker spatial separation and gender conditions for each group. Upper panels show results for specific gender conditions, and lower panels show results pooled for same-gender (MM and FF; gray color) and different gender conditions (FM and MF; blue color). **A**, **E**) NH listeners; **B**, **F**) bilateral HA users; **C**, **G**) bimodal CI+HA users; **D**, **H**) bilateral CI users. Error bars represent standard deviations around the mean.

A two-way repeated measures ANOVA was performed within each subject group with TMR thresholds as the dependent variable, and spatial separation (0° and ±60°) and gender difference (same and different) conditions as independent variables. Spatial separation between target and maskers was a significant factor for TMR improvement only in the NH group (p<0.001). Gender difference between target and maskers had a significant effect on TMR thresholds in NH, bilateral HA, and bimodal CI+HA groups (p<0.001 for all cases). A significant interaction between spatial separation and gender difference was observed only in the NH group (p<0.001). These trends are also apparent in Figure 3.

Only NH listeners benefited from spatial separation of the target and maskers. For NH listeners, mean TMR averaged across all target-masker pairs (mean and std = −9.9 ± 2.5 dB) for the spatially separated conditions was significantly greater than the mean TMR for the co-located conditions (mean and std = −1.1 ± 1.1 dB; F_1,20_ = 110.51, p<0.001, one-way ANOVA). This difference in TMR thresholds between co-located and spatially separated conditions indicates the amount of SRM, which shows how much speech recognition thresholds in noise are improved by spatial separation of the talker from the maskers. In contrast, for all of the HI groups, spatial separation did not improve the averaged TMR thresholds, meaning that no significant SRM was observed (bilateral HA: 1.1 ± 1.3 dB SRM; bimodal CI+HA: −0.7 ± 0.8 dB SRM; bilateral CI: 0 ± 0.3 dB SRM; p>0.3 for all HI groups).

Interestingly, in the NH group, mean SRM was significantly larger with same-gender maskers (11.7 ± 2.8 dB: difference between grey bars in Fig. 3E) than with different-gender maskers (6 ± 2.6 dB: difference between blue bars in Fig. 3E; F_1,20_ = 24.58, p<0.001). Bilateral HA users also had larger SRM with the same-gender masker (1.7 ± 1.8 and 0.4 ± 1.8 dB SRM for the same- and different-gender masker conditions, respectively), but the difference was marginally significant (F_1,22_ = 4.04, p=0.057). Neither CI group showed a significant difference in SRM between these two conditions (p>0.8).

Voice pitch differences between target and maskers, on the other hand, benefited NH, bilateral HA, and bimodal CI+HA listeners, but not bilateral CI listeners. This is especially apparent in the bottom row of Figure 3 which shows results pooled according to voice gender combination. TMR threshold improvements were observed from the same-gender to the different-gender conditions within each spatial separation configuration for NH listeners (Fig. 3E; difference between blue and gray bars), bilateral HA listeners (Fig. 3F) and bimodal CI+HA listeners (Fig. 3G). This difference in TMR thresholds between same-gender and different-gender conditions indicates the amount of VGRM, which indicates how much speech recognition thresholds in noise are improved by differences in gender between the target and masker.

The VGRM averaged across two target-masker spatial configurations was 5.7 ± 0.6 dB for NH listeners, 3.5 ± 1.4 dB for bilateral HA users, 3.4 ± 1.9 dB for bimodal CI+HA users, and 0.1± 0.2 dB for bilateral CI users. In particular, the VGRM for NH listeners was significantly larger in the co-located condition (8.5 ± 1.8 dB VGRM) than in the spatially separated condition (2.8 ± 1.1 dB VGRM, F_1,20_ = 77.91, p<0.001; Fig. 3E). A similar but not quite significant VGRM difference was also observed in bilateral HA users (co-location: 4.1 ± 1.9 dB VGRM; ±60° separation: 2.8 ± 1.4 dB VGRM, F_1,22_ = 5.55, p=0.052; Fig. 3F). The bimodal CI+HA users showed VGRM with almost no difference between the co-located and spatially separated conditions (co-location: 3.5 ± 2.1 dB VGRM; ±60° separation: 3.4 ± 1.9 dB VGRM; Fig. 3G). Bilateral CI users showed no VGRM for either spatial configuration (co-location: 0.1 ± 0.2 dB VGRM; ±60° separation: 0.1 ± 0.3 dB VGRM; Fig. 3G). Note that the amount of VGRM was much smaller for bilateral HA and bimodal CI+HA users compared to NH listeners in the co-located condition.

### Relationship of voice gender masking release to binaural pitch fusion range

The next step was to determine whether VGRM, the release from masking due to voice gender differences, is related to the width of binaural pitch fusion. Figure 4 shows individual VGRM with a fixed female target, i.e. the difference in TMR thresholds between the FF and FM conditions, plotted as a function of fusion range. Results are shown for the four listener groups in the co-located condition in the upper panels and the spatially separated condition in the lower panels. For NH listeners (Fig. 4A), the VGRM was negatively correlated with the fusion range only for the co-located target-masker condition (r=-0.698, p=0.017, two-tailed Pearson correlation test). In other words, listeners with narrow binaural pitch fusion ranges had larger VGRM - larger differences in TMR thresholds between same-gender (FF) and different-gender (FM) maskers - than did listeners with broad fusion. For both bilateral HA users and bimodal CI+HA users, a similar but stronger correlation between VGRM and binaural pitch fusion was observed in the co-located condition (bilateral HA: r=-0.873, p=0.001; bimodal CI+HA: r=- 0.862, p=0.001). Bilateral CI users showed no VGRM for either spatial configuration.

**Figure 4.**
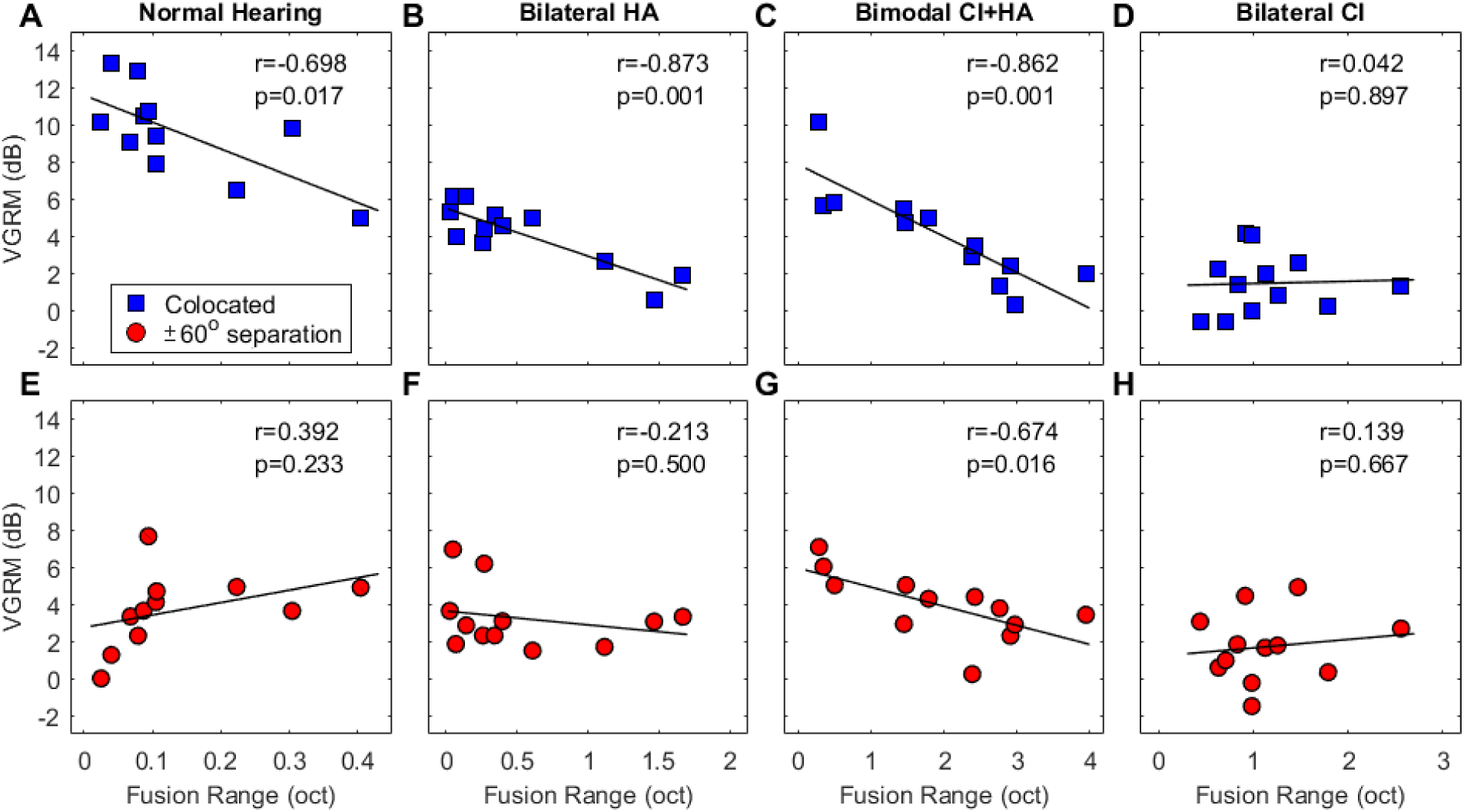
Correlations between binaural pitch fusion range and binaural voice gender release from masking (VGRM) for the fixed female target, i.e. the release in masking for the FM compared to the FF condition. Upper panels (**A-D**) show correlations for co-located conditions and lower panels (**E-H**) show correlations for the spatially separated conditions. **A**, **E**) NH listeners; **B**, **F**) bilateral HA users; **C**, **G**) bimodal CI+HA users; **D**, **H**) bilateral CI users.

Similar to the VGRM observed with the female target conditions, a significant negative correlation was also observed between VGRM with a male target and fusion range for NH listeners (r=-0.680, p=0.021), bilateral HA users (r=-0.774, p=0.003), and bimodal CI+HA users (r=-0.768, p=0.004) in only the co-located condition. Again, no VGRM was observed with a male target in bilateral CI users in either spatial configuration. All correlation results with different target-masker gender combinations are provided in Table 2. As hypothesized, only the same-versus different-gender comparisons (MM-MF, MM-FM, FF-MF, and FF-FM) showed statistically strong correlations between TMR differences and binaural pitch fusion in the co-located condition (see boldfaced correlation values in Table 2). It should be noted that a strong VGRM effect in the ±60° spatially separated configuration was observed only with the female target for the bimodal CI+HA users (FF-FM).

**Table 2.** Two-tailed Pearson correlation coefficients between binaural voice gender release from masking (VGRM) and binaural pitch fusion range widths for all listener groups in each spatial separation condition. Correlation values in bold face indicate significant results (*** p < 0.001; ** p < 0.01; * p < 0.05).

| Spatial separation | Target-masker gender combinations | Correlation r values (significance) |  |  |  |
| --- | --- | --- | --- | --- | --- |
|  |  | NH | Bilateral HA | Bimodal CI+HA | Bilateral CI |
| Co-location | FF-FM | **-0.698 (0.017)\*** | **-0.873 (<0.001)\*\*\*** | **-0.862 (<0.001)\*\*\*** | 0.042 (0.897) |
|  | FF-MF | -0.587 (0.058) | **-0.817 (<0.001)\*\*\*** | **-0.702 (0.011)\*** | 0.136 (0.672) |
|  | MM-MF | -0.513 (0.057) | **-0.806 (0.002)\*\*** | **-0.584 (0.046)\*** | 0.296 (0.350) |
|  | MM-FM | **-0.680(0.021)\*** | **-0.774 (0.003)\*\*** | **-0.768 (0.004)\*\*** | -0.115 (0.722) |
|  | MM-FF | -0.018 (0.958) | -0.338 (0.283) | -0.072 (0.824) | -0.167 (0.603) |
|  | MF-FM | -0.238 (0.481) | -0.148 (0.646) | -0.313 (0.174) | 0.135 (0.677) |
| ±60 degree separation | FF-FM | 0.392 (0.233) | -0.213 (0.500) | **-0.674 (0.016)\*** | 0.139 (0.667) |
|  | FF-MF | 0.341 (0.305) | -0.074 (0.820) | -0.392 (0.208) | 0.217 (0.498) |
|  | MM-MF | 0.202 (0.552) | -0.216 (0.500) | -0.290 (0.360) | 0.319 (0.313) |
|  | MM-FM | 0.336 (0.312) | -0.316 (0.317) | -0.476 (0.118) | 0.049 (0.879) |
|  | MM-FF | -0.186 (0.585) | -0.323 (0.306) | -0.028 (0.931) | -0.141 (0.661) |
|  | MF-FM | -0.026 (0.940) | -0.207 (0.519) | -0.263 (0.410) | 0.342 (0.276) |

In order to check that these correlations are not due to fusion range and VGRM both being correlated with overall poorer frequency resolution in one ear, a subset of NH and bilateral HA users were also tested on VGRM in a monaural listening condition. Figure 5 confirms that there are no correlations between binaural fusion range width and monaural VGRM in the female target (FF-FM) co-located condition in either NH or bilateral HA groups. It is interesting that in NH listeners, the reduced correlation is due primarily to improvement in VGRM in individual subjects with broad fusion in either the left or right monaural condition (rightmost symbols in Fig. 5A-B) compared to the binaural condition (rightmost symbols in Fig. 4A). This means that some NH listeners derive greater benefit from voice gender differences in the monaural listening conditions. Correlation results for all target-masker gender combinations are shown in Table 3.

**Figure 5.**
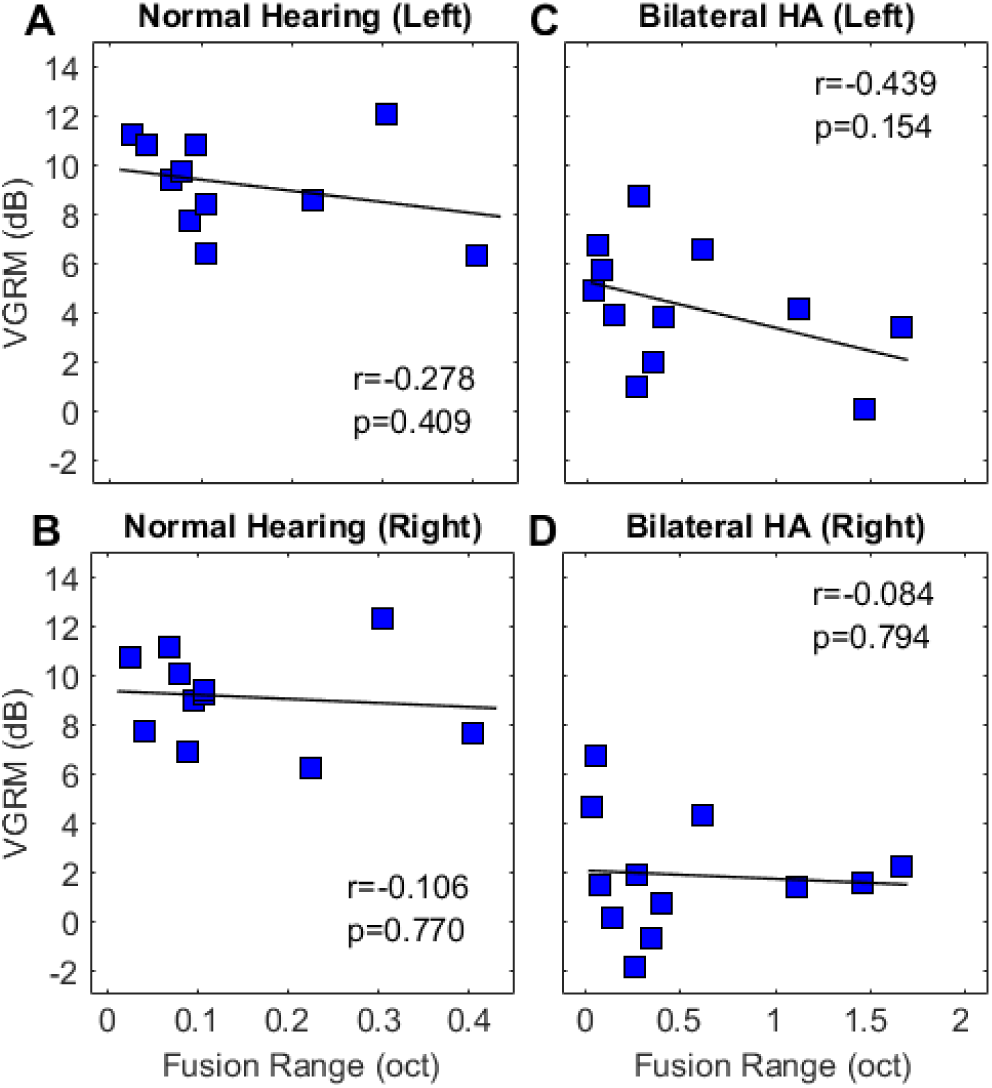
Correlations between binaural pitch fusion range and monaural voice gender release from masking (VGRM) for the fixed female target (FM-FF) in the co-located condition. Upper and lower panels show correlations for the left ear only and the right ear only conditions, respectively. **A**, **B**) NH listeners; **C**, **D**) bilateral HA users.

**Table 3.**
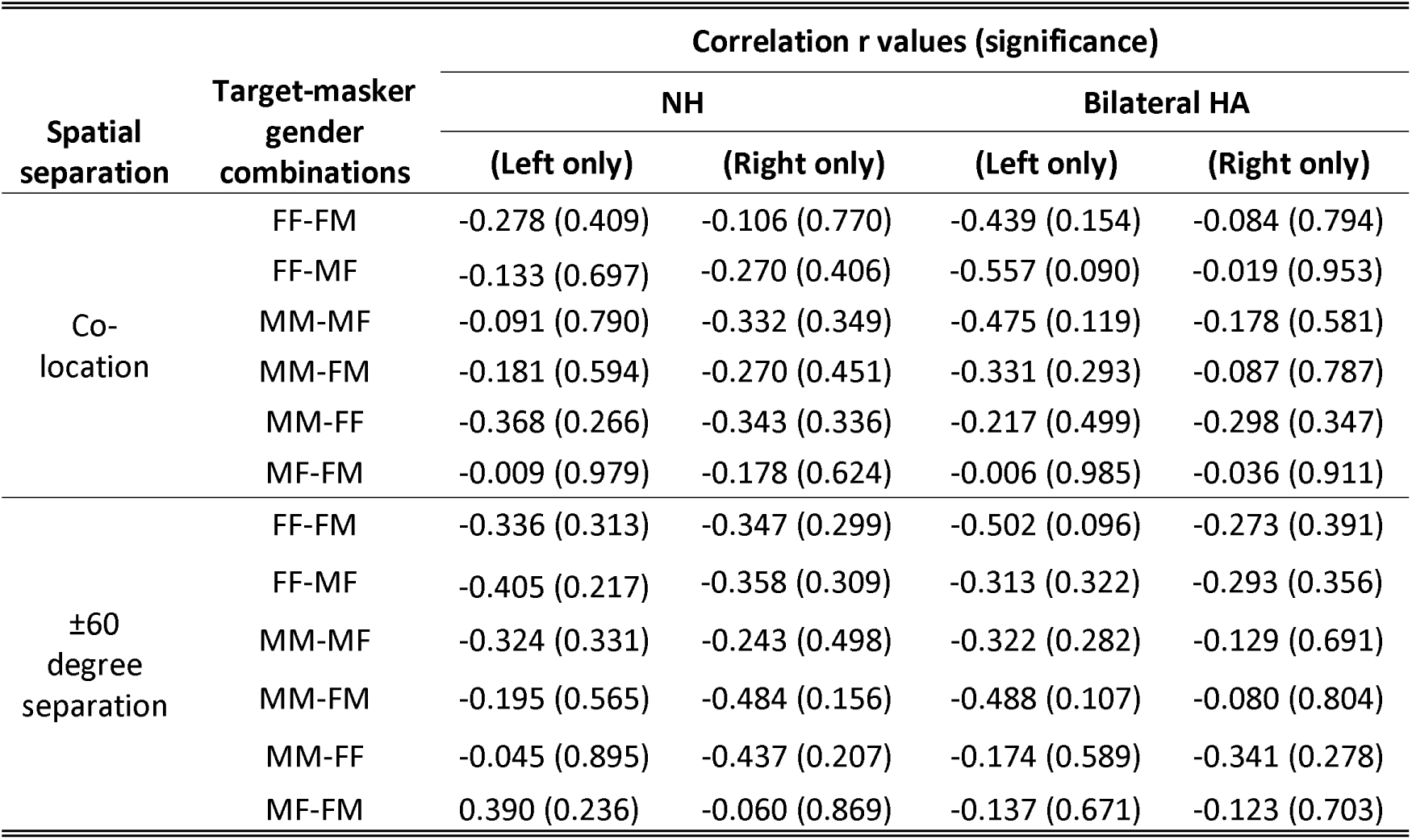
Two-tailed Pearson correlation coefficients between monaural voice gender release from masking (VGRM) and binaural pitch fusion range widths for NH and bilateral HA user groups in each spatial separation condition. Correlation values in bold face indicate significant results (*** p < 0.001; ** p < 0.01; * p < 0.05).

There were no significant correlations between VGRM and age in all listener groups. In HI groups, no significant correlations were also seen between VGRM and other subject factors such as age, average hearing thresholds in the reference and contralateral ear, age at onset of hearing loss, total duration of hearing loss, or duration of hearing device use. No significant correlations were seen between SRM and fusion range, or the subject factors described above in any listener groups.

## DISCUSSION

The ability to segregate a target talker from competing masker talkers is important for speech perception performance in multi-talker environments. Benefit from both spatial separation and voice pitch cues can be reduced in hearing-impaired listeners. These difficulties have previously been attributed to various factors (alone or in combination) on masking release performance: 1) a reduction in monaural temporal fine-structure sensitivity (Strelcyk and Dau, 2009; Neher et al., 2011; Summers et al., 2013), 2) a reduction in spectral or temporal modulation sensitivity (Bernstein et al., 2013), 3) a reduction in audiometric absolute thresholds and aging (Neher et al., 2011; Glyde et al., 2013; Besser et al., 2015; Gallun et al., 2013; Srinivasan et al., 2016; Jakien et al., 2017), and 4) a reduction in higher-order processing such as cognitive and linguistic abilities (Besser et al., 2015). To our knowledge, this is the first study to demonstrate an important role of another factor, abnormally broad binaural pitch fusion, in reduced binaural benefits for speech perception in multi-talker listening environments.

### Speech release from masking in NH and HI listeners

Performance in NH listeners for speech perception in competing talkers is consistent with previous literature showing benefits of both spatial separation and voice pitch differences between target and masker sources. Previous studies have also shown similarly greater benefit of spatial cues for same-gender conditions compared to different-gender conditions in both NH children and adults (Misurelli and Litovsky, 2012; 2015; Gallun et al., 2013; Gallun and Diedesch, 2013).

Unlike NH listeners, HI listeners did not show any spatial benefit. One likely reason for the reduced spatial benefit for HI users is that they have limited access to sound localization cues on the horizontal plane such as interaural time difference (ITD) and interaural level difference (ILD) cues. Previous studies have shown that ITD sensitivity is particularly important for localization performance and speech perception in noise (Gifford et al., 2013; 2014; Gallun and Diedesch, 2013; Swaminathan et al., 2016; Ellinger et al., 2017). Phase-locking and ITD sensitivity can both be impaired with hearing loss (Henry and Heinz, 2013; Dai et al., 2018). In addition, bilateral HA have reduced access and bilateral CI users have no access to ongoing ITD cues, because the hearing devices are not designed to coordinate their timing of stimulation of the auditory nerves across the ears (Brown et al., 2016). As for bimodal CI+HA users, their HA and CI devices are not linked. Thus, they do not communicate their processing schemes across the devices, such as compression ratio, which could alter ILDs (Byrne and Noble, 1998; Wiggins and Seeber, 2013). To our knowledge, it is less clear that ITD would be altered. To minimize any potential interaural cue distortion, the current study used symmetrical target-masker configurations (co-location and ± 60° separation) so that the image of both target and masker signals can appear in front, as opposed to the left or right due to reduced ILD, and all additional processing features for hearing devices were disabled to avoid altered ILD cues.

Note that in this study, effects of head shadow were also minimized due to the symmetrical target-masker configuration. In particular, spatial benefit in CI users can be observed in other spatial configurations which allow head shadow effects, such as asymmetrical target-masker configurations (Schleich et al., 2004; Bernstein et al., 2016) and directional microphones (van Hoesel and Tyler, 2003; Weissgerber et al., 2015).

Benefits of voice pitch differences were observed in both bilateral HA and bimodal CI+HA users, but not in bilateral CI users. For the bimodal CI+HA users, residual acoustic hearing in the HA ear allows the separation of different voice pitches due to better encoding of spectral information than in the CI ear. In contrast, bilateral CI users, who lack an acoustic hearing ear, have less access to this cue and have difficulty discriminating voice gender (Cleary and Pisoni, 2002; Fu et al., 2005; Gaudrain and Başkent, 2018). The way the CI processor is programmed does not provide sufficient spectral resolution to distinguish F_0_ and thus gender. The typical frequency-to-electrode allocation for the lowest frequency was 188-313 Hz in this study, which does not allow differentiation of the F_0_s for male (100 ± 7 Hz) and female talkers (204 ± 12 Hz) used in this study. The benefits observed for the bimodal CI+HA condition highlight the importance of retaining acoustic hearing when possible for maintaining voice discrimination and segregation abilities.

Neither age nor hearing loss was a major contributor to SRM or VGRM in any group, though age was barely correlated with SRM for NH listeners, similar to the previous study (Gallun et al., 2013). The lack of correlations with hearing loss in NH listeners could be due to a smaller sample size (N=11 versus N=34) and better hearing threshold criterion (<20 versus <50 dB HL) compared to the previous study.

### Correlations between binaural pitch fusion and voice gender release from masking

The present study shows that broad binaural fusion has a moderate to strong negative correlation with the ability to use voice pitch cues for speech perception with competing talkers; this is the first finding to investigate binaural fusion as a predictor. Previous studies have found that differences in age or hearing loss (alone or in combination) can explain some of the variance in the VGRM across subjects (Besser et al., 2015; Glyde et al., 2013). The proportion of variance accounted for by either factor was between 24% and 39% (R^2^ predictor, p<0.01). In this study, stronger negative correlations were observed between binaural fusion range and VGRM for NH listeners, bilateral HA users, and bimodal CI+HA users, especially in the co-located target-masker configuration. As reported in Table 2, the proportion of variance accounted for by binaural pitch fusion for VGRM was between 26% - 49% for NH listeners, 65% - 76% for bilateral HA users, and 34% - 74% for bimodal CI+HA users, higher than the amount of variance explained by age (R^2^ = 0.02, p<0.52; Glyde et al., 2013) or hearing loss (R^2^ = 0.39, p<0.001; Glyde et al., 2013) alone. Hence, broad binaural fusion is a stronger predictor for reduced VGRM than age or hearing loss.

In NH listeners, the lack of correlation between fusion range and SRM confirms that voice pitch separation ability between ears is not a major contributing factor in the ability to use spatial cues. Previously examined factors such as age, as well as binaural factors such as ITD sensitivity are likely to be more important in determining SRM.

Consistent with the correlation between fusion range and VGRM, some listeners with broad fusion reported fusing voices, and even reported the fusion of color words such as “green” and “red” into words not in the color set, such as “grain”. Spectral fusion of speech has been reported in the literature previously, but only for identical voice pitches in NH listeners (Cutting, 1976). This implies that broad fusion leads to abnormal spectral fusion and blending of words from voices of different pitches, just as pitch was observed to be averaged binaurally for tone stimuli (Reiss et al., 2014; Oh and Reiss, 2017). A visual analogue is the binocular averaging of color between the eyes (Hecht, 1928), in which information about the original colors is lost in fusion. Similarly, for speech, the fusion of words and sentences across multiple speakers would lead to the loss of information from the individual talkers at an early enough stage that the listener is unable to extract the original speech from any one stream.

The obligatory fusion and loss of information observed in hearing-impaired listeners suggest that some binaural spectral integration occurs before auditory object formation. This implies that abnormalities in binaural fusion due to hearing impairment may arise early in central auditory processing, such as at the brainstem level.

The correlation between binaural fusion and VGRM in bimodal CI+HA listeners is clinically important, because it explains a significant portion of the variability in benefit of bimodal hearing for speech perception in speech noise in the literature. More importantly, the reduced benefit in listeners with broad fusion reduces the overall average bimodal benefit across subjects and studies, so that bilateral implantation is still considered equivalent or superior to bimodal hearing by many clinicians. These findings demonstrate that clinical recommendations should be individualized based on binaural fusion; some bimodal CI+HA users with sharp fusion would lose as much as 10 dB of SNR benefit if they replaced the HA with a second CI. Bimodal CI+HA users with broad fusion would be more appropriate candidates for bilateral implantation.

In summary, based on these findings, the ideal scenario for hearing-impaired listeners is likely to be narrowly tuned fusion across ears similar to that experienced by NH listeners. Increased understanding of how abnormal binaural fusion affects binaural benefits for speech perception for hearing-impaired listeners is clinically essential for the future design of training- and device-based rehabilitative strategies to increase the benefits of binaural processing for speech perception in quiet and noise.

## ACKNOWLEDGEMENTS

We would like to thank Cochlear and MED-EL for providing equipment and software support for interfacing with the cochlear implant. This research was supported by grants R01 DC013307, P30 DC005983, and F32 DC016193 from the National Institutes of Deafness and Communication Disorders, National Institutes of Health.

